# Evolutionary dynamics of shared niche construction

**DOI:** 10.1101/002378

**Authors:** Philip Gerlee, Alexander R.A. Anderson

**Affiliations:** Integrated Mathematical Oncology, H. Lee Moffitt Cancer Center and Research Institute 12902 Magnolia Drive Tampa, FL 33612

## Abstract

Many species engage in niche construction that ultimately leads to an increase in the carrying capacity of the population. We have investigated how the specificity of this behaviour affects evolutionary dynamics using a set of coupled logistic equations, where the carrying capacity of each genotype consists of two components: an intrinsic part and a contribution from all genotypes present in the population. The relative contribution of the two components is controlled by a specificity parameter *γ*, and we show that the ability of a mutant to invade a resident population depends strongly on this parameter. When the carrying capacity is intrinsic, selection is almost exclusively for mutants with higher carrying capacity, while a shared carrying capacity yields selection purely on growth rate. This result has important implications for our understanding of niche construction, in particular the evolutionary dynamics of tumor growth.

---

In models of density-dependent growth the carrying capacity plays a pivotal role [1]. It represents the maximal size of the population, and is often viewed as an external limitation imposed by the environment on a growing population. Different genotypes might however experience differential carrying capacities by virtue of expressing different phenotypes, that vary in their degree of adaptation to survive at high densities [2]. This idea has for example been exploited in a game theoretical context [3], where it was assumed that the payoff received by each genotype influenced the carrying capacity, and not reproductive success, as is commonly assumed.

In many cases an increase in carrying capacity comes about through niche construction, whereby the organisms alter their environment in such a way that it can sustain a higher number of individuals [4]. For example the ant species *Myrmelachista schumanni* favours the growth of the tree *Duroia hirsuta*, in which it nests, by producing formic acid that is detrimental to other plants, and this in turn leads to more nesting sites for the ants [5]. Another example is the production of biofilm by certain bacterial species. This protective structure formed by polysaccharides is known to increase antibiotic resistance, but the chemical composition of the biofilm also influences colony size [6], and hence the carrying capacity of the strain. Lastly, we mention the importance of angiogenic factors released by tumor cells that stimulate the formation of new blood vessels, leading to increased nutrient and oxygen concentration, that facilitates higher cell densities and the growth of the tumour as a whole [7].

In all three examples there exists the possibility of cheating or free-riding on the genotype that facilitates the increased carrying capacity. The ability to do so largely depends on the specificity of the niche construction activity. If the modification of the niche is highly specific to the genotype that generates it, then most likely it is harder for other genotypes to exploit it, whereas a more general modification is easier to free-ride on. A natural question that arises in this context is how the specificity of the niche construction activity alters the evolutionary dynamics of the system.

A simple way of modelling and investigating this phenomena is by considering the carrying capacity of each genotype as being the sum of two components: an intrinsic carrying capacity and a contribution from other genotypes present in the ecosystem, the relative impact of the two components being controlled by a specificity parameter *γ*.

To investigate the effect of the niche construction specificity on the evolutionary dynamics we consider a multispecies system, where the number of organisms of type *i*, *x_i_* is governed by

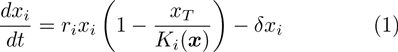

where *r_i_* > 0 is a species specific growth rate, 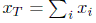 is the total population size, *K_i_*(***x***) is a species specific carrying capacity that depends on the current species composition ***x*** = (*x*_1_, *x*_2_,…,*x_n_*) and *δ >* 0 is a density independent death rate assumed equal for all species.

We assume that the carrying capacity for each species is determined both by a instrinsic/local and mean-field/global component and use a parameter *γ* ∈ [0, 1] to interpolate between the two. On one side of the spectrum, where specificity is maximal, we have the situation where each species only experiences its own intrinsic carrying capacity *k_i_*, and on the other side we have the case of minimal specificity, where all species experience the same carrying capacity *K*, that is given by the weighted mean of the constituent species carrying capacities, i.e.

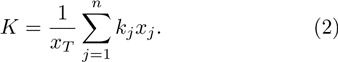

We now let

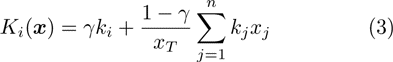

where *γ* = 1 corresponds to the local case (each species has a unique carrying capacity), and *γ* = 0 represents the global case where the carrying capacity is constant across all species. Please note that in the presence of a single species the system reduces to the standard logistic equation for all values of *γ*.

An important property of such a system is the stability of a resident population with respect to an invading mutant, and we now proceed to investigate this question, focusing on the impact of the degree of specificity *γ*.

To this end we consider the situation where only *n* = 2 species are present, in which case the system of equations simplifies to

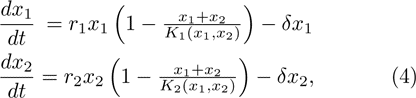

where

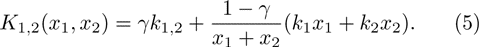

The steady-state, corresponding to a monomorphic population, is given by ***x****\** = (*x*_1_*, x*_2_) = (*k*_1_(1 − *δ/r*_1_), 0), where we have assumed (without loss of generality) that species 1 is the resident and 2 is the mutant.

The mutant can invade the resident population if the steady-state is unstable, i.e. if at least one of the eigenvalues of the Jacobian *J* evaluated at ***x****\** has a positive real part.

The Jacobian at ***x****\** is given by

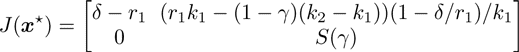

where

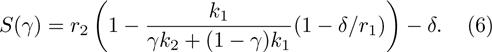

The eigenvalues of *J* evaluated at the monomorphic steady-state ***x****\** are therefore given by *λ*_1_ = *δ − r*_1_ and *S*(*γ*). Now *δ − r*_1_ *<* 0 (or else the equilibrium concentration of the resident would be negative), and hence the ability of a mutant to invade depends on the sign of *S*(*γ*), where *S >* 0 corresponds to the ability of the mutant to invade.

We start by investigating the extreme points *γ* = 1 and 0, that correspond to a locally versus globally determined carrying capacity. For the case with maximal specificity we have

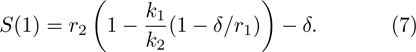

For a small death rate (*δ ≪ r*_1_) we have *S*(1) ≈ *r*_2_(1 − *k*_1_/*k*_2_), which implies that *S >* 0 if and only if *k*_2_ *> k*_1_, i.e. the mutant can invade if it has a larger carrying capacity than the resident.

In the case of minimal specificity (*γ* = 0) we have

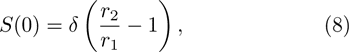

which implies that the mutant can invade if *δ >* 0 and *r*_2_ *> r*_1_.

This means that when the carrying capacity is genotype specific, there is selection for mutants with a higher carrying capacity, while the case in which the carrying capacity is global and determined by all genotypes, there is selection for mutants with a higher growth rate. In the latter case a non-zero death rate is also required for any mutant to invade.

In order to get a better understanding of the impact of the specificity we plot the curve *S*(*k*_2_*, r*_2_) = 0 in the (*r, k*)-parameter space for three different choices of *γ* (see figure 1). The regions above (and to the right) of the curves correspond to the subset of mutant characteristics, in terms of the relative growth rate (*r*_2_/*r*_1_) and relative carrying capacity (*k*_2_/*k*_1_), for which a mutant can invade. In the global case (*γ* = 0) the ability to invade depends only on the mutant growth rate whereas in the local case (*γ* = 1) dependence is almost exclusively in the *k*-direction. As is evident from the intermediate case (*γ* = 0.5), the transition between the two extreme cases occurs for low values of *γ*, and sharpness is controlled by the death rate *δ*. For smaller values of *δ* the transition is even more sudden.

**FIG. 1.**
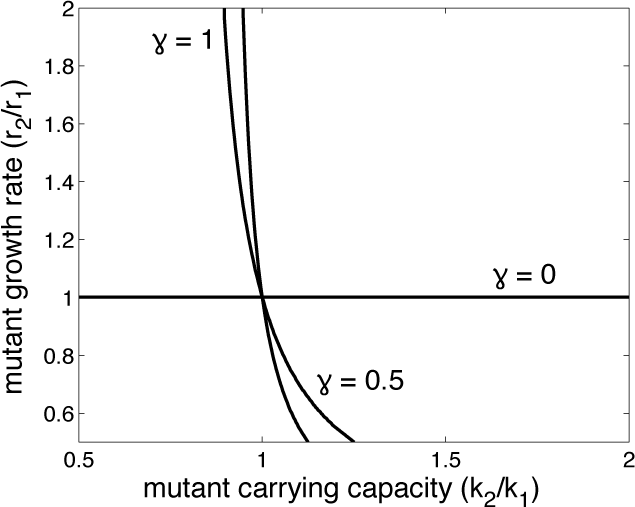
Invasion as a function of mutant *r* and *k*. The region above (and to the right) of the curves correspond to the subset of mutant characteristics, in terms of the relative growth rate (*r*_2_/*r*_1_) and relative carrying capacity (*k*_2_/*k*_1_), for which a mutant can invade. In the local case (*γ* = 1) dependence is almost exclusively in the *k*-direction, whereas in the global case (*γ* = 0) only the mutant growth rate affects the ability to invade. As is evident from the curve corresponding to *γ* = 0.5, the transition between the two extreme cases occurs for small values of *γ*. The death rate was set to *δ* = *r*_1_/10.

The impact of the death rate *δ* on the transition from the local to the global dynamics is further examined in figure 2. Panel A shows the minimal carrying capacity *k_min_* required for a mutant with *r*_2_/*r*_1_ = 1.1 to invade a resident population, while B shows the minimal mutant growth rate required when the ratio of the carrying capacities is fixed at *k*_2_/*k*_1_ = 2. From the two plots it is evident that the mutants carrying capacity largely determines its ability invade, even for small values of *γ*, when the benefit from niche construction is relatively unspecific. However, we note that the sharpness of the transition decreases as the death rate approaches the resident growth rate.

**FIG. 2.**
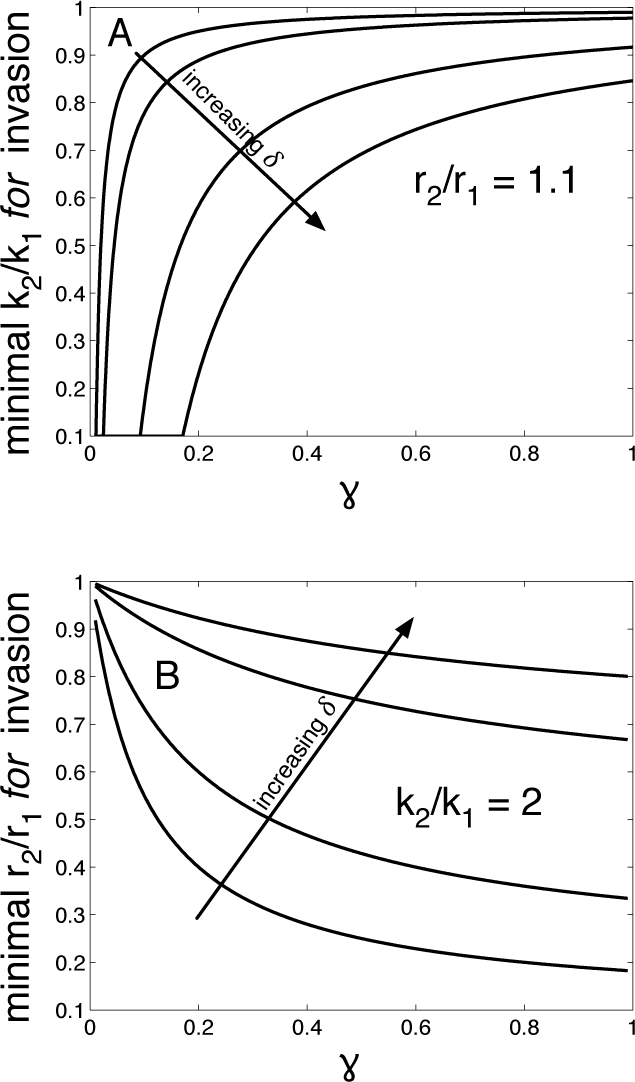
Ability to invade as a function of *γ*. (A) shows the minimal carrying capacity required for a mutant to invade when the ratio between mutant and resident growth rate is *r*_2_/*r*_1_ = 1.1. (B) shows the minimal mutant growth rate required for invasion when the ratio between mutant and resident carrying capacity is *k*_2_/*k*_1_ = 2. The different lines correspond to death rates equal to *δ* = *r*_1_ × (0.1, 0.2, 0.5, 0.67).

These results suggest that a biological system governed by (1) could experience different evolutionary outcomes depending on the value of the specificity parameter. From our analysis of the two-species case it seems as if a system evolving under *γ* = 1, would, because of the selection on carrying capacity, increase its total population size *x_T_* at a faster rate than the same system would for *γ* < 1. In addition we would expect evolution to favour lower growth rates as the specificity *γ* increases.

In order to test these hypotheses we extended the model to take into account evolutionary dynamics. We consider the coupled system of equations (1) with a single wild-type present with *r*_0_ = 1, *k*_0_ = 10^6^ and *x*_0_(0) = 1. The system is solved numerically with time step Δ*t* = 0.01, and at each time step we introduce a new mutant with probability *µx_T_* Δ*t*, where *µ* = 10*^−^*^7^ is the per capita mutation rate. The growth rate and carrying capacity of the mutants are for simplicity assumed to be independent of the resident population and drawn at random from the intervals *r_m_* ∈ [0, 2*r*_0_] and *k_m_* ∈ [0, 2*k*_0_]. This is of course a gross simplification, since one would expect correlations both between the resident and mutant phenotype, and between *r_m_* and *k_m_* [8], but sufficient for our purposes.

Figure 3A shows an example of the evolutionary dynamics of (1) when *γ* = 0. Initially the population is invaded by two mutants that increase the total population size, but eventually a mutant appears, that, while successful at displacing the resident, lowers the population size. Figure 3B shows the dynamics for *γ* = 1, from which we can discern a clear trend towards a selection of a higher carrying capacity, where each mutant that successfully invades the population displaces the resident and reaches fixation at a higher population size than the previous resident. Figure 3C shows the total population size *x_T_* after *t* = 1000 time units as a function of *γ*, and the inset shows the mean growth rate 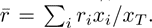 The results are averaged over 500 simulations per value of *γ* and confirms our two hypotheses – the total population size is an increasing function of *γ*, while the average growth rate decreases (roughly linearly) with *γ*.

**FIG. 3.**
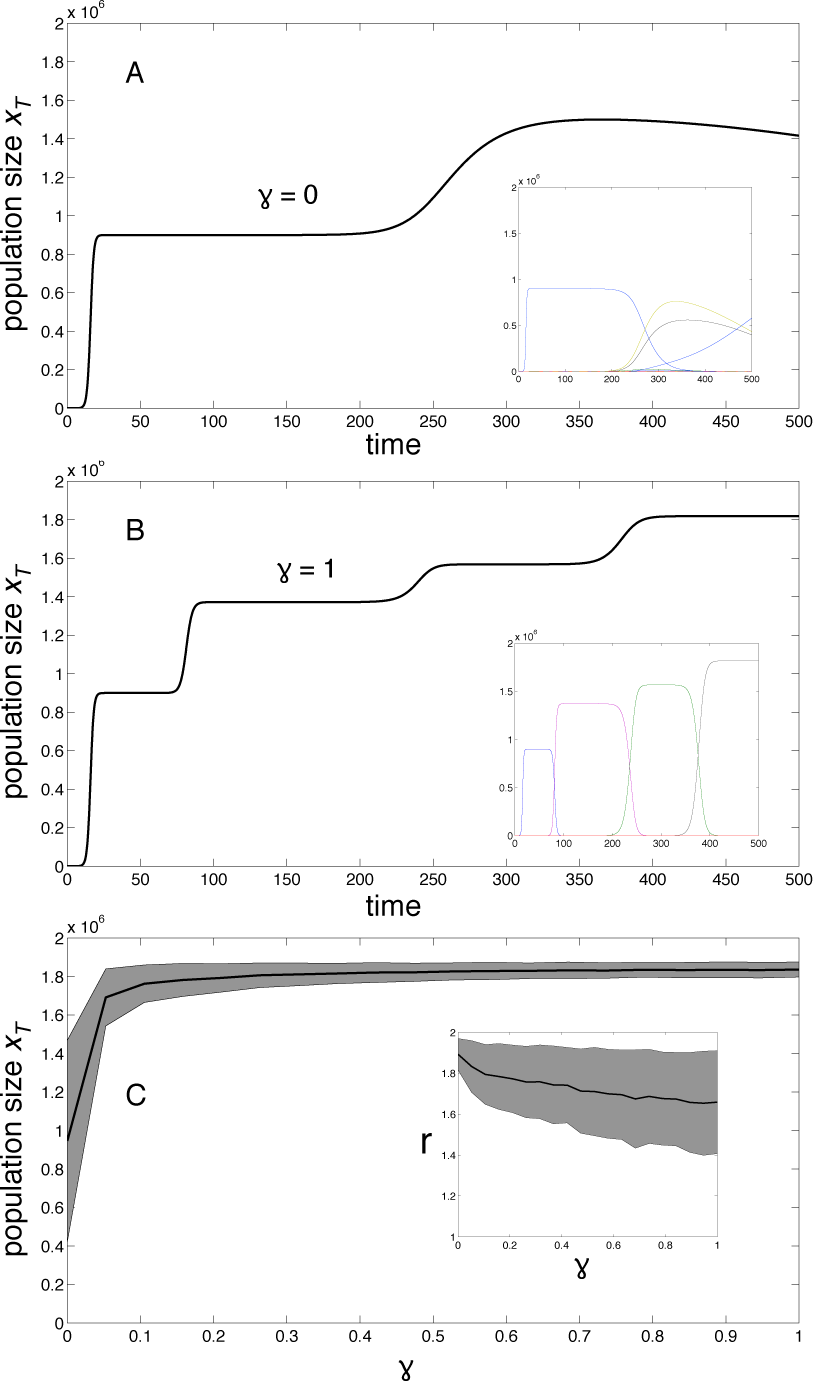
Evolutionary dynamics for different values of the specificity parameter *γ*. (A) shows a simulation in which *γ* = 0, where the total population size first increases, but a successful invasion of a clone with a smaller carrying capacity leads to a decrease in population size. (B) shows a simulation for *γ* = 1, and as expected from the analysis of the two-species system, clones with higher carrying capacity are selected for, which leads to an increase in the total population size. (C) shows the total population size after *t* = 1000 time steps as a function of *γ*, and the inset shows the resulting average growth rate. The results are averaged over 500 simulations for each value of *γ* (shaded area shows one standard deviation) and confirms the trend seen in the above panels – the total population size is an increasing function of *γ*, whereas the average growth rate decrases.

These results can be understood from an intuitive point of view by considering the dynamics at the two extremes of niche construction specificity. When *γ* = 0 all genotypes contribute in proportion to their abundance to a carrying capacity shared by all genotypes. The situation therefore resembles that of a public goods game in which all participants receive benefit from the public good. In analogy with the well-known result that the public goods game can be invaded by free-riders that contribute less than the resident, our system can be invaded by mutants with a smaller carrying capacity. However, this mutant needs to have a growth rate that exceeds that of the resident in order to spread in the population. In the case of maximal specificity (*γ* = 1) a single mutant, when introduced into a resident population at carrying capacity, can only spread if, when a resident dies, it has a chance of dividing and replacing the former resident. Since the total population size is at the carrying capacity of the resident, the mutant will only divide and spread if its carrying capacity is higher than the residents.

Moving between the two extremes, by varying the specificity parameter *γ*, results in a rather sharp transition between the two regimes (see fig. 2). Even for small values of *γ* the ability of a mutant to invade is dominated by its carrying capacity, e.g. a mutant with *r*_2_/*r*_1_ = 1.1 needs a carrying capacity of *k*_2_ ≈ *k*_1_ already for *γ* = 0.5. This is also reflected in the evolutionary model, where the total population size seems to reach a plateau value already for small values of *γ* (see fig. 3C). This suggests that under the dynamics proposed in this Letter the long-term dynamics of a population are more likely to resemble that of fig. 3B rather than A, i.e. the total population size is expected to increase with the introduction of each mutant.

The relationship between selection for carrying capacity and growth rate, usually termed *r/K*-selection has a long history in evolutionary biology dating back to the seminal work of MacArthur in the 1960s [9]. Although the paradigm has largely been replaced by more detailed studies of life history evolution, it has provided important insight into the evolutionary dynamics of density-dependent selection. For example it has been shown that populations that evolve in stable and mild environments will experience selection for high *K*-values, while populations in harsh seasonal environments will evolve towards higher *r* [2]. The results presented here suggest a novel mechanism by which selection can favour either an increased carrying capacity or growth rate. However, the value of *γ* can itself be viewed as an outcome of selection, most likely influenced by the cost of maintaining a specific response.

The dependence on *γ* has important implications for our understanding of tumor growth. Naively one would expect that since many factors (e.g. angiogenic and autocrine signalling) that positively impact tumor growth are diffusible and hence highly unspecific in their impact on different subclones, they would be subject to exploitation. This would then lead to a collapse of the population, but our results suggest that there is a persistent selection towards a higher carrying capacity even for very low levels of specificity. In addition, the results provide a suggestion as to why tumor tissue has a much higher density than normal tissue – consecutive selective sweeps of clones with higher carrying capacity have occurred during disease progression.

Differential carrying capacities have in fact largely been disregarded in models of tumor growth, which have instead focused on increased growth rates and adaptation to adverse environmental conditions (low nutrients, cytotoxic therapy etc.). The results presented herein underline the importance of taking carrying capacity into account when attempting to explain tumour growth dynamics, and highlight the potential prognostic value of increased tissue density.

The model that we have analysed disregards a number of aspects related to density dependent selection and niche construction. For example we have not considered the cost involved in increasing the carrying capacity, and also assumed that inter- and intraspecific competition are equal. Despite this the model provides novel insight into the impact of niche construction specificity on evolutionary dynamics, and hints at the importance of including differential carrying capacities when considering density regulated growth.

## References

[1] D. MonteLuna, B. Brook, M. ZetinaRejón, and V. CruzEscalona, Global ecology and biogeography 13, 485 (2004).

[2] J. Roughgarden, Ecology, 453 (1971).

[3] S. Novak, K. Chatterjee, and M. A. Nowak, Journal of theoretical biology 334, 26 (2013).

[4] D. Erwin, Trends in Ecology & Evolution 23, 304 (2008).

[5] M. Frederickson and D. Gordon, Proceedings of the Royal Society B: Biological Sciences 274, 1117 (2007).

[6] I. Ramos, L. Dietrich, A. Price-Whelan, and D. Newman, Research in microbiology 161, 187 (2010).

[7] P. Hahnfeldt, D. Panigrahy, J. Folkman, and L. Hlatky, Cancer Research 59, 4770 (1999).

[8] L. Mueller, Annual Review of Ecology and Systematics, 269 (1997).

[9] R. MacArthur, Proceedings of the National Academy of Sciences of the United States of America 48, 1893 (1962).

